# Pregnancy success in mice requires appropriate cannabinoid receptor signaling for primary decidua formation

**DOI:** 10.1101/2020.08.04.236869

**Authors:** Yingju Li, Amanda Dewar, Yeonsun Kim, Sudhansu K. Dey, Xiaofei Sun

## Abstract

An early event after implantation is stromal cell transformation to decidual cells (decidualization) that support embryo development. In mice, this process begins at the antimesometrial (AM) pole with differentiation of stromal cells into epithelial-like cells (epithelioid cells) surrounding the implantation chamber. This is an avascular zone called the primary decidual zone (PDZ), and considered to function as a transient, size-dependent permeable barrier to protect the embryo from maternal circulating harmful agents, including immunoglobulins, immune cells, microorganisms and other noxious agents. This zone forms on day 5 afternoon and becomes fully established on day 6 with the loss of the crypt epithelium. The PDZ gradually degenerates with the appearance of the secondary decidual zone (SDZ) around the PDZ that peaks on day 8 in mice. Decidualization is critical for early pregnancy in mice and humans. We show that cannabinoid/endocannabinoid signaling influences decidualization in early pregnancy. Mice deficient in two major cannabinoid receptors, CB1 and CB2, show compromised PDZ. We found that angiogenic factors are dysregulated in *Cnr1*^*-/-*^*Cnr2*^*-/-*^ mice with defective PDZs, resulting in the abnormal presence of blood vessels and macrophages in this zone; disruption of the PDZ compromises pregnancy outcomes. Using an *in vitro* decidualization model, we found that *Cnr1* levels increase in mouse stromal cells and human uterine fibroblast (Huf) cells undergoing decidualization and that limiting CB1 signaling in these cell types suppresses decidualization *in vitro*. Since endothelial cells express *Cnr2* and decidual cells express *Cnr1*, we hypothesize that angiogenic events driven by CB2 are integrated with CB1 in decidual cells, leading to proper PDZ formation, a critical step for pregnancy success.

## Introduction

Decidualization is a critical pregnancy event that follows embryo implantation. During the initial stages of decidualization in mice, stromal cells transform into epithelioid cells (epithelial-like) surrounding the implanting embryo and form an avascular zone. This region is called the primary decidual zone (PDZ) and is critical to embryo development and successful pregnancy outcomes in mice (1). The PDZ encircles the implantation chamber (crypt) on day 5 of pregnancy and is fully established by day 6. Because the PDZ is avascular, this zone is devoid of maternal immune cells, which protects embryos from circulating maternal insults (2). Recently we have shown that Scribble (Scrib), a scaffold protein and a component of the planar cell polarity (PCP), plays a key role in PDZ formation (3). However, molecular regulation in the formation of the PDZ is still far from clear.

Natural cannabinoids (tetrahydrocannabinol, the most potent psychoactive component) and endocannabinoids (anandamide and 2-arachidonoilglycerol) signal through two membrane receptors, CB1 (encoded by *Cnr1*) and CB2 (encoded by *Cnr*2). Both CB1 and CB2 are G-protein coupled receptors in the Gi/o and Gq families (4). CB1 is mainly expressed in the central nervous system (CNS), but also appears in peripheral tissues including the heart, testis, liver, small intestine and uterus. In contrast, CB2 is mostly expressed in immune cells (5). We have recently shown that CB2 is also expressed in uterine endothelial cells (6). Deletion of *Cnr2* increases uterine edema before implantation and causes suboptimal implantation (6).

Our work has previously shown that elevated cannabinoid signaling is detrimental to early pregnancy events in mice (7). Furthermore, we have shown that *Cnr1*^*-/-*^*Cnr2*^*-/-*^ mice have compromised implantation (6), suggesting that silencing of endocannabinoid signaling is also detrimental to early pregnancy events. In the study, we showed that the litter sizes of double mutant mice, but not single mutant mice, are substantially smaller than wild-type (WT) mice. We also found that endothelial cells with *Cnr2* deletion play an important role in embryo implantation (6). Interestingly, resorption rate is lower in *Cnr2*^*-/-*^ females than in *Cnr1*^*-/-*^*Cnr2*^*-/-*^ females during the midgestational stage, indicating a critical role for CB1 in early pregnancy. Since CB1 abundance is quite low in the uterus, its role in early pregnancy remains obscure.

In this study, we investigated the roles of endocannabinoid signaling in decidualization using mouse models with suppressed CB1 and CB2 and a primary culture of *Cnr1*^*-/-*^ stromal cells. The results show that double mutant mice have compromised PDZ formation, resulting in suboptimal pregnancy outcomes. Further investigation revealed that *Cnr1* is expressed in decidual cells. These results, combined with our previous knowledge of CB2 expression in endothelial cells, show for the first time that angiogenic actions incited by CB2 are integrated by CB1 in decidual cells to form the avascular PDZ, which is critical for pregnancy success.

## Results

### *Cnr1*^*-/-*^*Cnr2*^*-/-*^ females have suboptimal decidualization

Previously we showed that *Cnr1*^*-/-*^*Cnr2*^*-/-*^ uteri have defective decidualization and consequent increases in midgestational resorption rates. However, the mechanism by which the absence of CB1 and CB2 signaling compromises uterine decidualization remains unclear. To explore this further, WT and *Cnr1*^*-/-*^*Cnr2*^*-/-*^ females were mated with WT males. Induction of *Ptgs2*, a rate-limiting enzyme for prostaglandin formation, at the crypt (implantation chamber) epithelium and underlying stroma at the site of implantation has been shown to be critical to initiate decidualization (8). Thus, we examined the expression of *Ptgs2* on day 5 of pregnancy to capture the influence of *Ptgs2*. The expression pattern and signal intensity of *Ptgs2* is comparable in WT and *Cnr1*^*-/-*^*Cnr2*^*-/-*^ females (Fig 1a), indicating stimulation from embryo implantation is well received by *Cnr1*^*-/-*^*Cnr2*^*-/-*^ uteri. However, the *in situ* hybridization of *Bmp2*, a morphogen expressed by cells undergoing decidualization at the site of embryo implantation (9), reveals that stromal cell decidualization is defective in *Cnr1*^*-/-*^*Cnr2*^*-/-*^ females on day 5 of pregnancy (Fig 1b). Expression of *Hoxa10*, also critical for decidualization, shows a similar reduction in *Cnr1*^*-/-*^*Cnr2*^*-/-*^ implantation sites (Supplemental fig 1). Decidual defects in *Cnr1*^*-/-*^*Cnr2*^*-/-*^ uteri continue on day 6 of pregnancy. The tridimensional (3D) structure of implantation sites reveals that, on day 6 of pregnancy, decidual responses in WT mice promote the uterine lumen to form an arch shape, whereas in *Cnr1*^*-/-*^*Cnr2*^*-/-*^ implantation sites, the lumen remains flat (Fig 2a). The domain of *Bmp2* positive signals is reduced in *Cnr1*^*-/-*^*Cnr2*^*-/-*^ implantation sites (Fig 2b). In contrast, stromal cells after decidual transformation downregulate the expression of *Bmp2*. More decidualized cells around the implantation chamber negative for *Bmp2* signals are observed on day 6 of pregnancy in WT mice as compared to *Cnr1*^*-/-*^*Cnr2*^*-/-*^ implantation sites (Fig 2b). With decidualization in progress, *Ptgs2* expression transitions from the antimesometrial side of the crypt to the mesometrial side in the WT crypt on day 6 of pregnancy. In *Cnr1*^*-/-*^*Cnr2*^*-/-*^ implantation sites, cells on the lateral side of the implantation chamber show *Ptgs2* signals, suggesting impeded progress of decidualization (Fig 2c).

**Figure 1.**
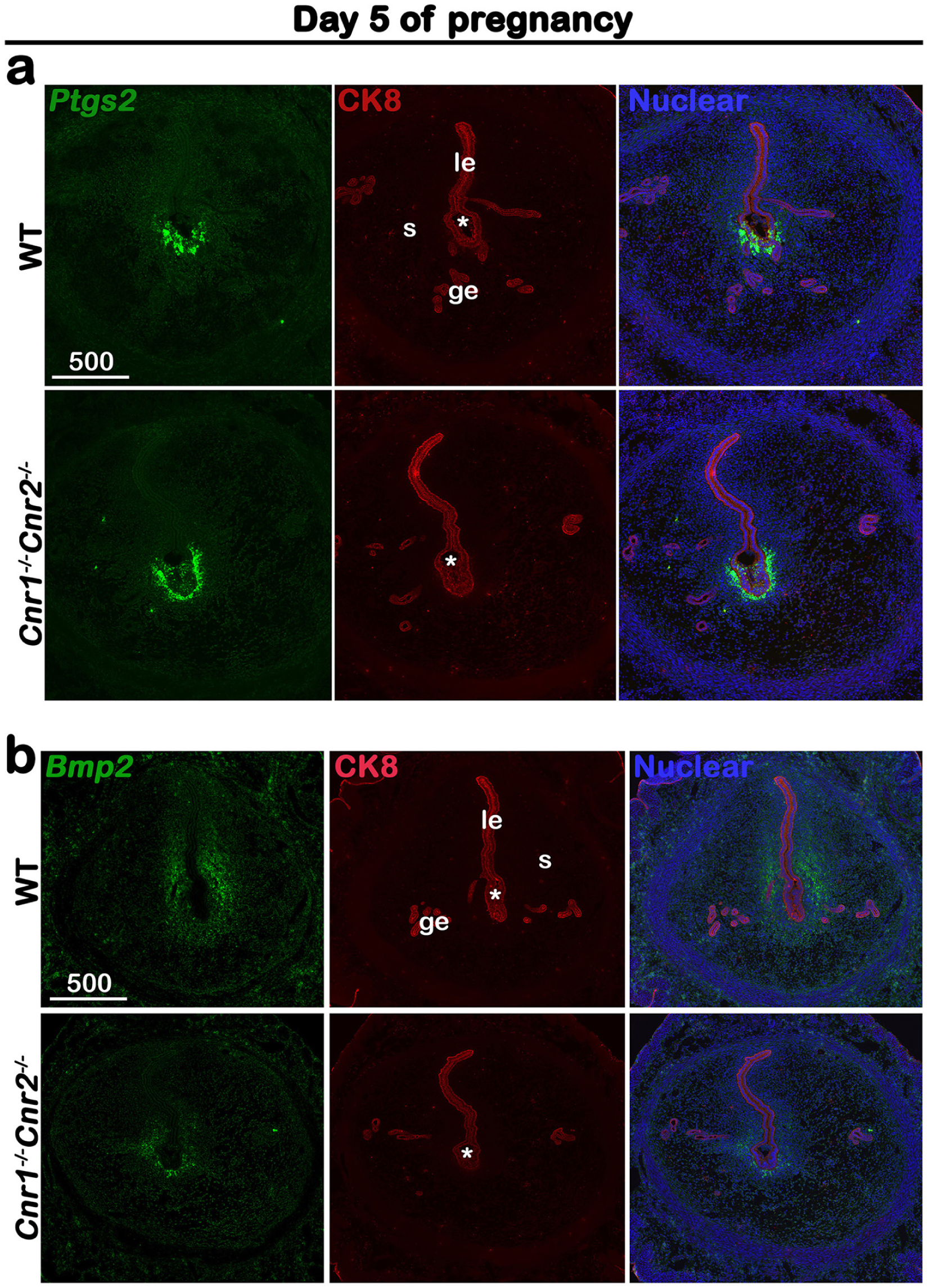
*Cnr1*^*-/-*^*Cnr2*^*-/-*^ females show normal responses to embryonic stimulation during implantation, but decidual process is compromised. (a) *In situ* hybridization of *Ptgs2* in uteri on day 5 of pregnancy. The expression patterns of *Ptgs2* critical for implantation show no significant difference in WT and *Cnr1*^*-/-*^*Cnr2*^*-/-*^ females. CK8 staining outlines uterine epithelial cells. (b) *In situ* hybridization of *Bmp2* in uteri on day 5 of pregnancy. Decidual responses in *Cnr1*^*-/-*^*Cnr2*^*-/-*^ females are much weaker than those in WT females. le, luminal epithelium; s, stroma; ge, glandular epithelium; Asterisks, positions of embryos; Scale bars, 500 μm. All images are representative of three independent experiments.

**Figure 2.**
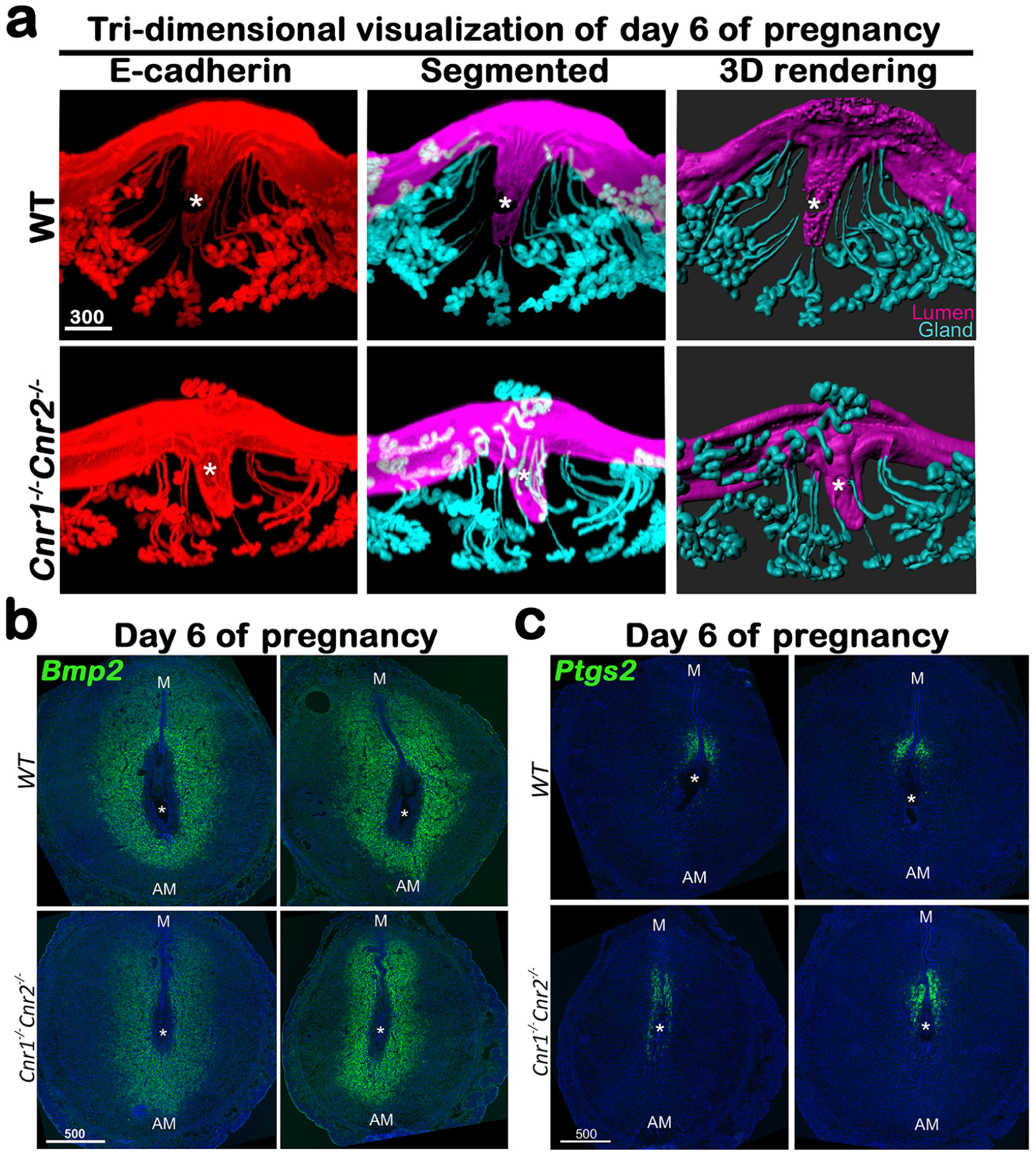
*Cnr1*^*-/-*^*Cnr2*^*-/-*^ females have suboptimal decidualization. (a) 3D visualization of day 6 implantation sites in WT and *Cnr1*^*-/-*^*Cnr2*^*-/-*^ females. Images of E-cadherin immunostaining, segmented, and 3D rendered images of day 6 implantation sites in each genotype show compromised decidual responses in *Cnr1*^*-/-*^*Cnr2*^*-/-*^ females. Scale bars, 300 μm. (b and c) *In situ* hybridization of *Bmp2* and *Ptgs2* in uteri on day 6 of pregnancy. M, mesometrial; AM, anti-mesometrial; Asterisks, positions of embryos; Scale bars, 500 μm. All images are representative of three independent experiments.

### Macrophages are retained around the implantation chambers of compromised deciduae in *Cnr1*^*-/-*^*Cnr2*^*-/-*^ females

Hematopoietic cells, including macrophages, are distributed throughout mouse uteri prior to implantation, though these cells are not present in the decidual zone after implantation (10,11). We observed that the distribution of hematopoietic cells, as revealed by a pan hematopoietic marker CD45, is comparable in *Cnr1*^*-/-*^*Cnr2*^*-/-*^ and WT uteri on day 4 of pregnancy (Fig 3a). This observation is further confirmed by the distribution of macrophages, which are demarcated by F4/80 staining (Fig 3b). These results suggest that hematopoietic cells normally enter the uterus of *Cnr1*^*-/-*^*Cnr2*^*-/-*^ mice before implantation. On day 6 of pregnancy, while most macrophages are absent from the WT implantation chamber, some macrophages remain around the *Cnr1*^*-/-*^*Cnr2*^*-/-*^ embryos, albeit reduced in number (Fig 3c). The significant lingering of macrophages in the *Cnr1*^*-/-*^ *Cnr2*^*-/-*^ endometrium (Fig 3d) suggests that impeded decidual responses in *Cnr1*^*-/-*^*Cnr2*^*-/-*^ females fail to dispel macrophages efficiently. Collectively, the result suggests that the normally developed decidua protects the developing blastocyst from inflammatory insult from macrophages and that this safe-guard mechanism is compromised in double-mutant mice.

**Figure 3.**
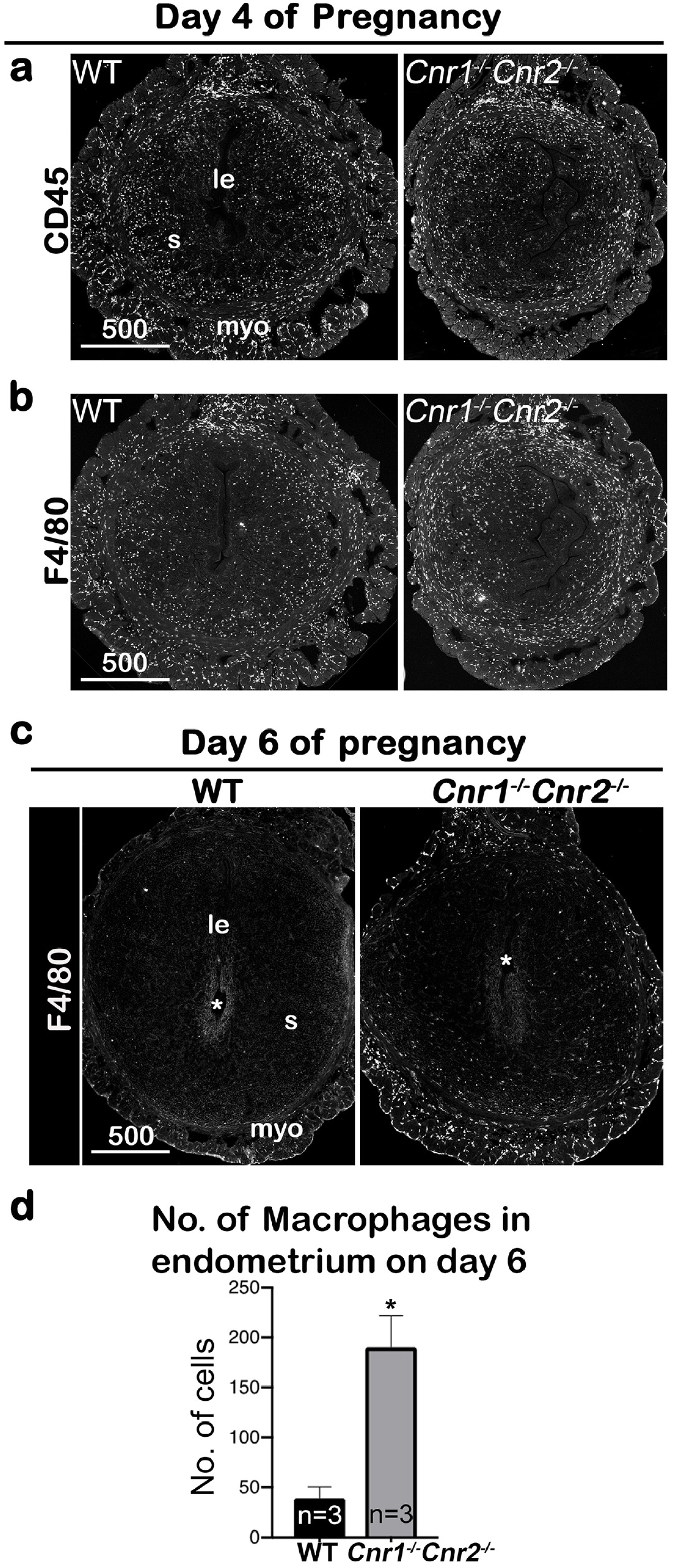
Macrophages are retained around implantation chambers of compromised deciduae in *Cnr1*^*-/-*^*Cnr2*^*-/-*^ females. (a) Uterine leukocytes in pre-implantation are highlighted by CD45, which is expressed on all leukocytes. (b) Immunofluorescent staining of F4/80 on day 4 of pregnancy to mark macrophages. (c) Immunofluorescent staining of F4/80 on day 6 of pregnancy. (d) The numbers of macrophages within the endometrial domains are quantified using 3 sections obtained from 3 different animals in each genotype. le, luminal epithelium; s, stroma; myo, myometrium; Asterisks, positions of embryos; Scale bars, 500 μm. All images are representative of three independent experiments.

### The formation of the primary decidual zone is compromised in *Cnr1*^*-/-*^*Cnr2*^*-/-*^ implantation sites

On day 6 of pregnancy, the first stratum of the decidual zone, called the primary decidual zone (PDZ), is fully established. As previously stated, the PDZ is an avascular zone comprised of stromal cells undergoing epithelial-like transformation and is impermeable to maternal immune cells; as such, this zone is indispensable to protect and support further embryo development (2). The finding that macrophages remain close to *Cnr1*^*-/-*^ *Cnr2*^*-/-*^ implantation chambers prompted us to examine the formation of the PDZ on day 6 of pregnancy. PDZ formation requires Scrib signaling (3), yet the Scrib positive zone domain is much reduced in *Cnr1*^*-/-*^*Cnr2*^*-/-*^ females (Fig 4a). This result is also supported by the presence of blood vessels, revealed by immunostaining of FLK1, a reliable marker of blood vessels, in the *Cnr1*^*-/-*^*Cnr2*^*-/-*^ PDZ on day 6 of pregnancy (Fig 4b). The presence of blood vessels in *Cnr1*^*-/-*^*Cnr2*^*-/-*^ PDZ is significantly abundant than that in WT PDZ (Supplemental fig 2). *Vegfa*, which promotes angiogenesis by targeting its receptor FLK1 (12), shows a dynamic spatiotemporal expression pattern in periimplantation uteri (13). *Vegfa* expression is downregulated in WT PDZ on day 6 of pregnancy, whereas positive *Vegfa* signals are observed in the PDZ of *Cnr1*^*-/-*^*Cnr2*^*-/-*^ mice (Fig 4b), similar to FLK1 positive cells in the same area. These results indicate that normal disappearance of blood vessels is hindered in *Cnr1*^*-/-*^*Cnr2*^*-/-*^ PDZ, accompanied by lingering macrophages around the embryo.

**Figure 4.**
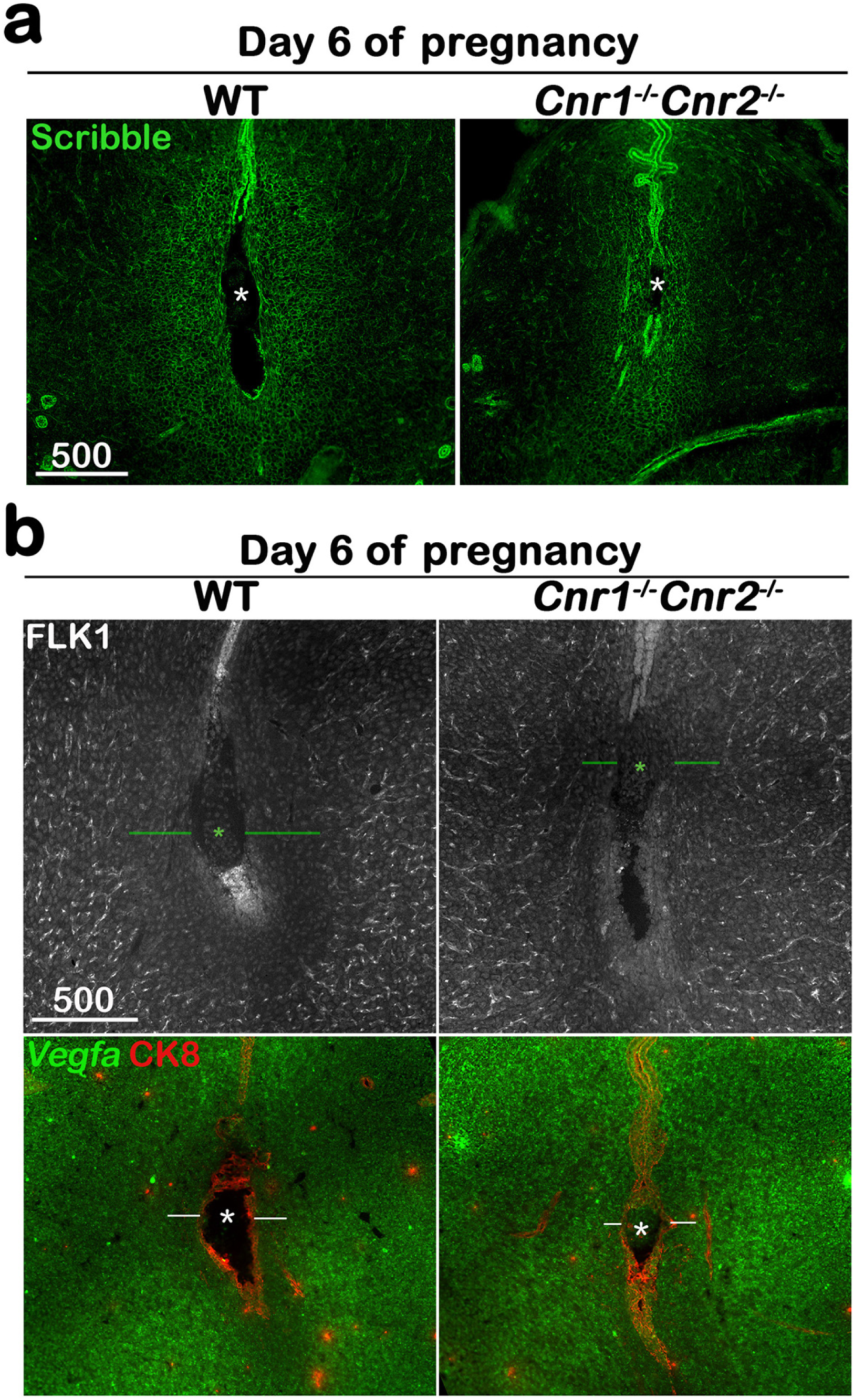
The formation of the primary decidual zone is compromised in *Cnr1*^*-/-*^*Cnr2*^*-/-*^ implantation sites. (a) The primary decidual zone is highlighted by Scribble on day 6 of pregnancy. (b) Immunofluorescence staining of FLK1 and *in situ* hybridization of *Vegfa* in day 6 pregnant uteri. Blood vessels and higher *Vegfa* signals are observed in the PDZ area (indicated by green or white lines) of *Cnr1*^*-/-*^*Cnr2*^*-/-*^ implantation sites. CK8 staining outlines epithelial cells. Asterisks, positions of embryos; Scale bars, 500 μm. All images are representative of three independent experiments.

### Retention of blood vessels in *Cnr1*^*-/-*^*Cnr2*^*-/-*^ PDZ is associated with multiple angiogenic factors

Proangiogenic angiopoietin-1 (ANGPT1), targeting TIE2 receptors on endothelia, is critical to assemble newly formed vasculature and stabilize the vascular network and integrity (14,15). In contrast, angiopoietin-2 (ANGPT2), an ANGPT1 antagonist (16), promotes endothelial cell apoptosis and vascular regression. *Angpt1* is expressed in most stromal/decidual cells but is absent in the PDZ region of day 6 implantation sites in WT mice (Fig 5a). However, *Angpt1* expression remains in the PDZ of *Cnr1*^*-/-*^*Cnr2*^*-/-*^ mice, which facilitates the stability of vessels in the region (Fig 5a). The signal intensity of *Angpt1* is quantified in Supplemental figure 3. In contrast to *Angpt1, Angpt2* is expressed in decidualized stromal cells surrounding the implantation chamber at the mesometrial side in WT females (Fig 5b). The domain expressing *Angpt2* and the signal intensity are much reduced in *Cnr1*^*-/-*^*Cnr2*^*-/-*^ implantation sites (Figure 5b and Supplemental fig 3).

**Figure 5.**
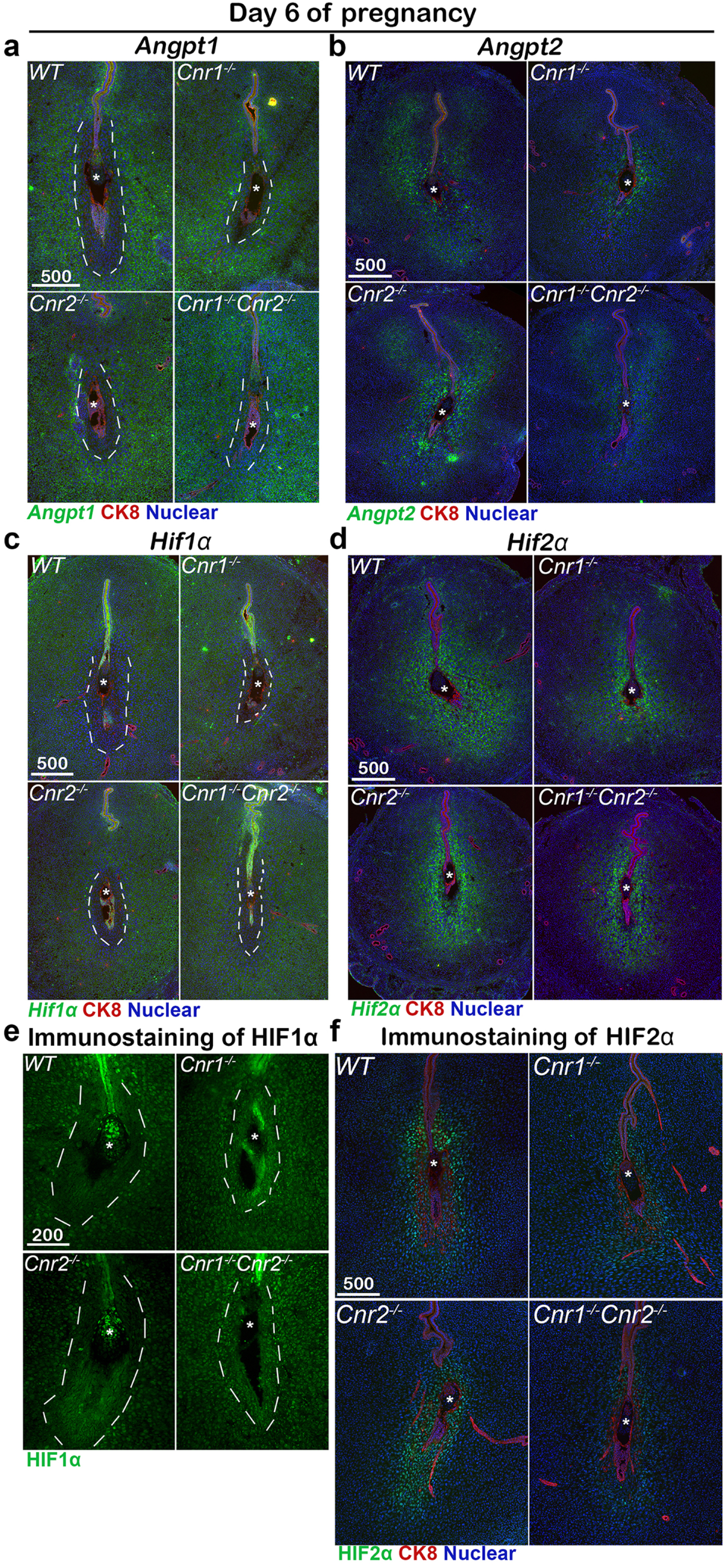
Angiogenic factors are misregulated in *Cnr1*^*-/-*^*Cnr2*^*-/-*^ and *Cnr1*^*-/-*^ PDZ. (a-d) *In situ* hybridization of *Angpt1, Angpt2, Hif1α* and *Hif2α* on day 6 of pregnancy. CK8 staining outlines uterine epithelial cells. (e and f) Immunostaining of HIF1*α* and HIF2*α* on day 6 of pregnancy. Dotted lines outline the PDZs. Asterisks, positions of embryos; Scale bars, 500 μm. All images are representative of three independent experiments.

These results indicate that opposing functions between ANGPT1 and ANGPT2 at implantation sites maintain a vascular network that supports normal implantation; in *Cnr1*^*-/-*^*Cnr2*^*-/-*^ implantation sites, these opposing functions are dysregulated. To further determine the roles of CB1 and CB2 in the regulation of ANGPTs, RNA levels of *Angpts* were examined in *Cnr1*^*-/-*^ and *Cnr2*^*-/-*^ mice. Intriguingly, the expression pattern of *Angpt1* and *Angpt2* in *Cnr1*^*-/-*^ implantation sites recapitulate their patterns in *Cnr1*^*-/-*^*Cnr2*^*-/-*^ counterparts (Fig 5a and 5b), suggesting that defects in ANGPT regulation are due to CB1 deficiency.

Hypoxia-inducible factors (HIFs) are transcriptional regulators of angiopoietins and vascular endothelial growth factors (VEGFs) (17,18). Interestingly, we found the expression pattern of *Hif1α* is similar to that of *Angpt1*, showing broad expression in stromal cells but lacking expression in WT PDZs on day 6 of pregnancy (Fig 5c). On the other hand, the pattern of *Hif2α* expression is similar to that of *Angpt2* with positive signals in stromal cells close to implantation sites (Fig 5d). PDZs in *Cnr1*^*-/-*^ and *Cnr1*^*-/-*^*Cnr2*^*-/-*^ mice show increased *Hif1*α expression, as well as decreased *Hif2α* expression compared with WT females on day 6 of pregnancy (Fig 5c and d, Supplemental fig 3). The mRNA expression patterns of *Hif1*α and *Hif2α* were further confirmed by immunostaining using antibodies specific to *Hif1*α and *Hif2α* (Fig 5e and 5f). These results suggest that increased *Angpt1* expression with decreased *Angpt2* expression in the PDZs of *Cnr1*^*-/-*^ *Cnr2*^*-/-*^ stabilizes blood vessels in this region. The differential expression patterns of *Hif1*α and *Hif2α* indicate that *Angpt1* and *Angpt2* are perhaps differentially regulated by HIF1α and HIF2α, respectively. The similar patterns of angiogenic factors in *Cnr1*^*-/-*^ and *Cnr1*^*-/-*^ *Cnr2*^*-/-*^ mice suggest that CB1 plays a key role in PDZ formation.

### *Cnr1* is induced in mouse and human decidual cells

CB2 is primarily expressed in immune cells. Using a *Cnr2* reporter mouse line (19), we have recently shown that uterine endothelial cells express CB2 (6). Studies show that deletion of *Cnr1* in the mouse uterus compromises pregnancy outcomes (20), even though *Cnr1* levels are quite low in non-pregnant uterine cells. We observed that *Cnr1* levels increase in implantation sites from day 4 to day 8 (the peak of decidual response) (Fig 6a). Since most uterine epithelial cells are removed in implantation sites on day 8, we attribute the increase to changes in stromal/decidual cells. We isolated and purified uterine stromal cells on day 4 of pregnancy and induced them to decidualize *in vitro* by hormone treatments according to our established protocol (21). Levels of *Cnr1* increased about 3-fold in WT and *Cnr2*^*-/-*^ decidualizing cells as compared to non-decidualized stromal cells in each genotype (Fig 6b).

**Figure 6.**
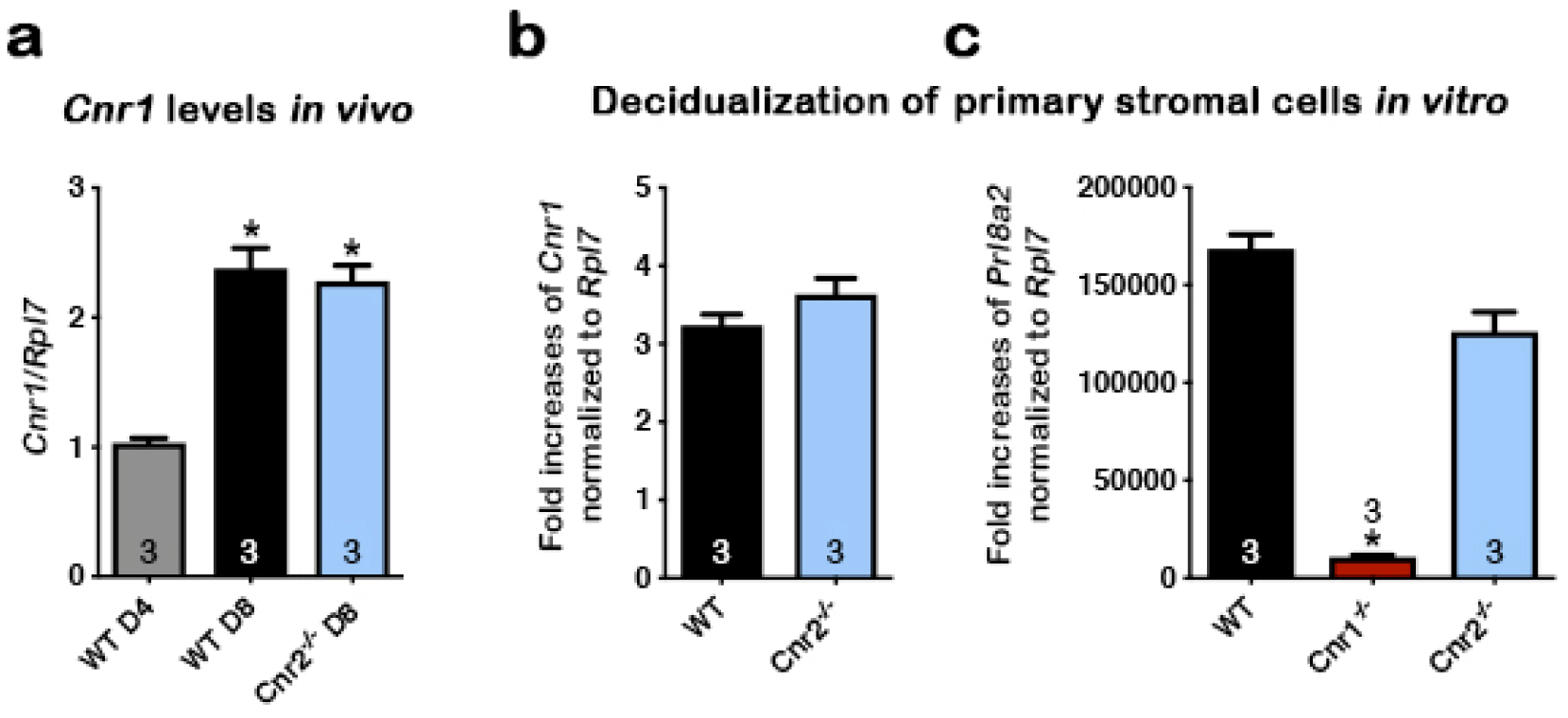
CB1 expression in mouse uterine stromal cells increases during decidualization. (a) Levels of *Cnr1* are higher at the peak of decidual response on day 8 of pregnancy as compared to day 4 of pregnancy. (b) *Cnr1* is induced in primary culture of WT and *Cnr2*^*-/-*^ stromal cells undergoing decidualization *in vitro*. (c) Decidualization is severely reduced in primary *Cnr1*^*-/-*^ stromal cells after decidual stimulation *in vitro. Prl8a2* expression served as indicator of decidual responses. Values are mean ± SEM; Numbers on bars represent sample sizes; *P< 0.05, *Student’s t* tests.

To study roles of *Cnr1* during decidualization, stromal cells collected from both *Cnr1*^*-/-*^ and WT uteri were induced to a decidual reaction *in vitro. Prl8a2*, a marker gene expressed in mouse decidual cells, was significantly lower in *Cnr1*^*-/-*^, but not *Cnr2*^*-/-*^, decidual cells (Fig 6c), suggesting deletion of *Cnr1* in stromal cells compromises decidualization. Interestingly, the induction of *CNR1* is even more robust in human uterine fibroblast (Huf) cells when decidualized *in vitro*. A steady increase of *CNR1* was observed during the cell transformation, and eventually *CNR1* increased by more than 100-fold compared with cells before decidualization (Fig 7a). The decidual responses in Huf cells in the 6 days after induction were confirmed by a rise in two decidual markers, *PROLACTIN* (*PRL*) and *IGFBP1* (Fig 7b and c). The morphology of Huf cells changed from a spindle shape to more of a round shape (Fig 7d and Supplemental fig 4), which was also reflected by increases in nuclear circularity during decidualization (Fig 7e and 7f). Cell circularity is calculated as a ratio of the minor axis versus the major axis (Fig 7e). The nuclear circularity in decidualized Huf cells is significantly higher than in cells before decidualization (Fig 7f). Although *CNR1* levels were 100-fold higher in decidualized cells compared to control cells (Fig 7a), *CNR2* levels show no significant changes (Supplemental fig 5), suggesting CB1 plays a more important role in human decidualization.

**Figure 7.**
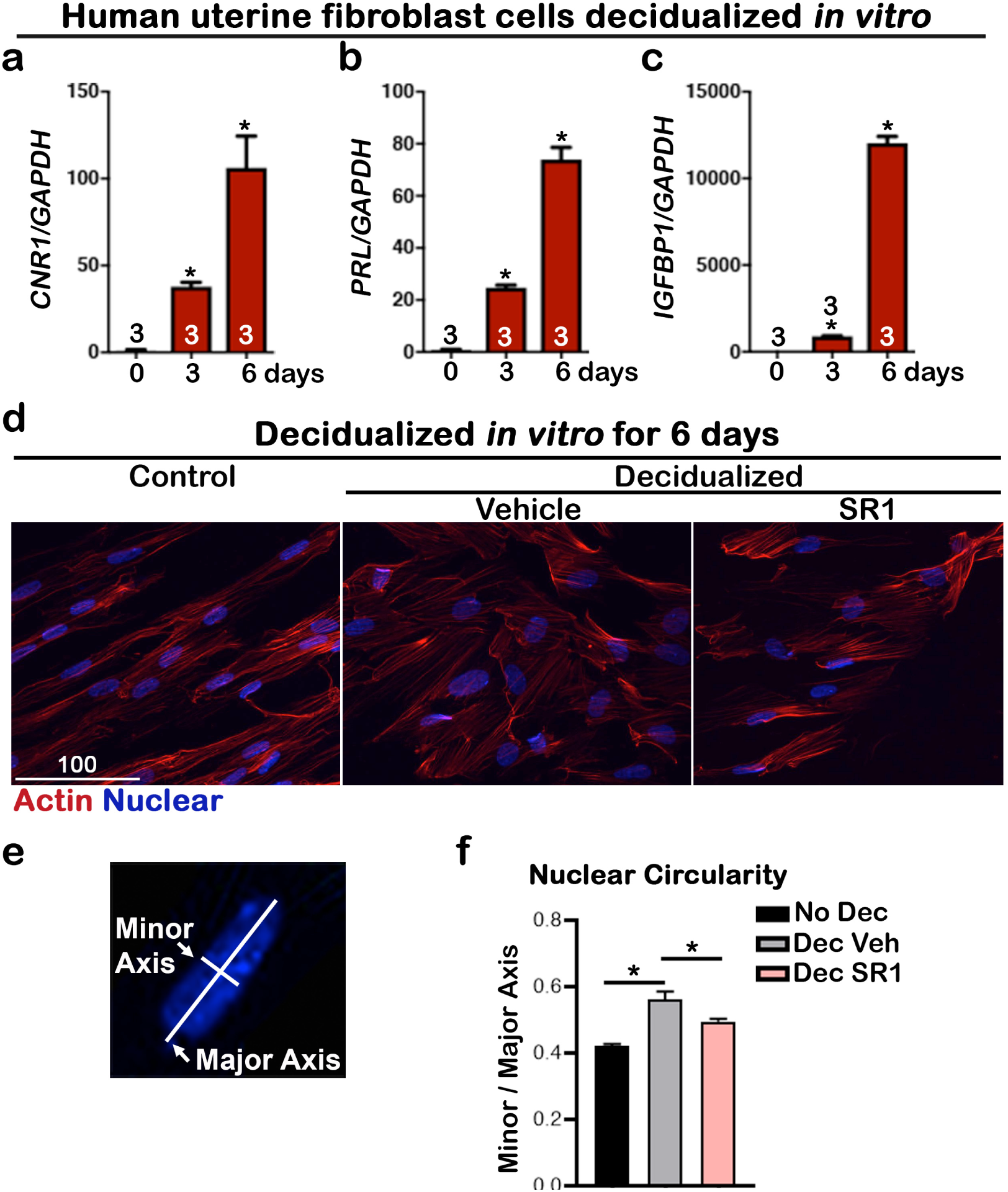
*CNR1* levels are upregulated in human uterine fibroblasts during *in vitro* decidualization. (a) Levels of *CNR1* gradually increase in Huf cells undergoing decidualization. (b and c) Decidual responses are marked by significant increases in *PROLACTIN* and *IGFBP1* levels in Huf cells. (d) Morphology of Huf cells changes from spindle shapes to more round shape; SR1 treatment compromises the morphological change in Huf cells. Cells are outlined with Actin staining. (e) A scheme depicts the measurement of minor and major axes for calculating nuclear circularity. (f) Nuclear circularity of Huf cells before and after decidualization. SR1 treatment compromises the decidual process. Six fields were randomly chosen in 3 different culture wells. Values are mean ± SEM; Numbers on bars are sample sizes; *P< 0.05, *Student’s t* tests.

To further study the biological significance of the induction of *CNR1*, Huf cells were treated with a specific CB1 antagonist, SR141716 (SR1, 2 µM), during decidualization. The decidual process in SR1-treated Huf cells was significantly compromised, as revealed by changes in *PRL* and *IGFBP1* levels (Fig 8a and 8b), two established decidual markers. The levels of *VEGFA, ANGPT1, ANGPT2* and *HIF2α* were significantly downregulated by SR1 treatment during decidualization (Fig 8), suggesting induction of *CNR1* during decidualization is critical to this process. Taken together, the upregulation of *Cnr1* in decidual cells in both mice and humans plays a key role during decidual cell transformation as supported by both *in vivo* and *in vitro* studies.

**Figure 8.**
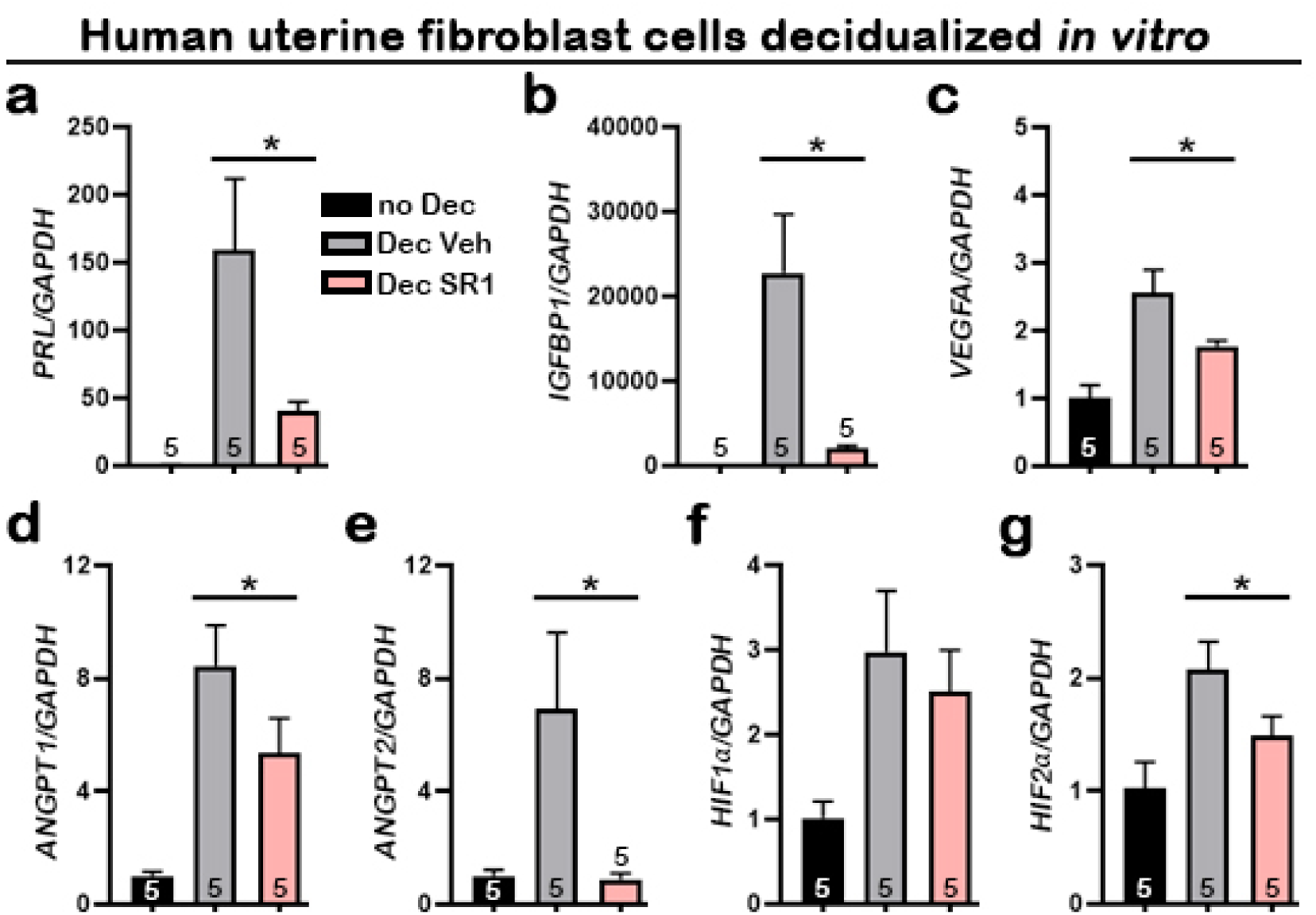
Suppression of CB1 impairs normal Huf cell decidualization and expression of angiogenic factors. (a-g) qPCR of *PRL, IGFBP1, VEGFA, ANGPT1, ANGPT2, HIF1α* and *HIF2α* using RNA collected 3 days after decidualization. Values are mean ± SEM; Numbers on bars are sample sizes; *P< 0.05, *Student’s t* tests.

## Discussion

In this study, we show that a complex interplay is orchestrated by cannabinoid signaling in decidualization by forming the PDZ along with changes in angiogenic features in the decidual bed. In *Cnr1*^*-/-*^*Cnr2*^*-/-*^ females, this organization is compromised due to defective PDZ formation and dysregulated angiogenic homeostasis. PDZ is avascular in WT implantation sites, whereas blood vessels are present in *Cnr1*^*-/-*^*Cnr2*^*-/-*^ PDZ, accompanied by an increased presence of macrophages.

Angiogenesis and vascular remodeling are hallmarks of implantation, decidualization and placentation (13,22,23). Retention of blood vessels in *Cnr1*^*-/-*^*Cnr2*^*-/-*^ PDZ is associated with misregulated angiogenic factors: *Angpt1*, which stabilizes vasculature, is abnormally expressed in *Cnr1*^*-/-*^*Cnr2*^*-/-*^ implantation sites. *Angpt2*, which destabilizes preexisting vasculature for *Angpt1* sprouting (16), is downregulated in *Cnr1*^*-/-*^*Cnr2*^*-/-*^ implantation sites. Our results also show that *Hif1α* has a similar expression pattern to *Angpt1* in WT PDZ on day 6 of pregnancy, while *Hif2α* expression pattern overlaps with that of *Angpt2*. This suggests that deletion of both *Cnr1* and *Cnr2* impacts PDZ formation and establishment of decidualization through regulation of angiogenic and transcriptional factors. The levels of *Cnr1*, which are usually low in stromal cells of non-pregnant females, increase during the decidual process in both humans and mice. Using an *in vitro* decidualization model, we further showed that decidualization is compromised using *Cnr1*^*-/-*^ stromal cells. Consequently, defective PDZ formation results in more resorption later in pregnancy in *Cnr1*^*-/-*^*Cnr2*^*-/-*^ females.

Most previous work examining the roles of endocannabinoids in decidualization applied cannabinoids to stromal cells. Although some pharmacological studies using CB1 antagonists suggest CB1 mediates the effects of endocannabinoids on stromal cells, there is no genetic evidence to support this conclusion. Using stromal cells isolated from pregnant rats, a previous report showed that an endocannabinoid anandamide (AEA) interferes with stromal differentiation (24). *In vivo* application of AEA in pseudopregnant rats impaired decidualization with reduced levels of Cox1 and VEGF (24). There was evidence for a similar observation in humans. In this respect, AEA impairs cell proliferation and differentiation in the human endometrial stromal cell line and primary cultures of decidual fibroblasts from term pregnancy (25). There is further evidence that oxidative metabolites of AEA by Cox2 may compromise the normal decidual process (26,27).

In contrast to the ectopic application of endocannabinoids, our study used mouse models with suppressed cannabinoid signaling. The studies cited above seem to contradict our current findings that cannabinoid signaling is detrimental to decidualization in rats and humans. However, we have shown previously that either amplified or silenced cannabinoid signaling can interfere with early pregnancy (7). These results suggest that optimal endocannabinoid signaling contributes to pregnancy success, reinforcing our conclusion that normal decidualization requires appropriate cannabinoid signaling.

The function of ANGPT2 in angiogenesis largely depends on the presence of VEGF (28,29) in that VEGF, ANGPT1 and ANGPT2 function cooperatively to promote angiogenesis (30,31). ANGPT2 destabilizes the preexisting vasculature, which consequently responds to angiogenic stimuli. The activated vasculature is extended and stabilized by ANGPT1 (29). In the absence of VEGF, ANGPT2 destabilizes vasculature, which undergoes vascular regression. In implantation sites, VEGF and ANGPT1 have similar expression patterns on day 6 of pregnancy (13), but neither is present in the PDZ; only ANGPT2 is expressed in this zone. The consequent avascular zone in the PDZ is considered conducive to the implanting embryo. In the current study, *Angpt1* expression is retained in *Cnr1*^*-/-*^*Cnr2*^*-/-*^ PDZ with reduced expression of *Angpt2*, so that vasculature and macrophages are still observed in *Cnr1*^*-/-*^*Cnr2*^*-/-*^ PDZs.

Hypoxia inducible factors (HIFs) are key modulators of the transcriptional responses to hypoxic stress. HIFs are heterodimers of an α-subunit and a stable β-subunit (ARNT). Of the 3 isoforms, HIF1α and HIF2α are the most studied and structurally similar. HIF1α is considered to have a broader expression pattern in most cells, whereas HIF2α is selectively induced in certain cell types, including endothelial cells and renal interstitial cells (32). In our study, *Hif1α* is expressed mostly in stromal cells except for the PDZ cells of day 6 implantation sites, while *Hif2α* is induced in PDZ. The transcriptional regulation of HIFs on angiopoietins is specific to tissue and cell-type. For example, in primary human endothelial cells, HIF1α increases mRNA levels of *ANGPT2* and *ANGPT4* but not *ANGPT1* (33). In the current study, the overlapping expression of *Hif1α* and *Angpt1*, in tandem with the similar expression pattern of *Hif2α* and *Angpt2*, suggests that HIF1α increases *Angpt1* expression in stromal cells and HIF2α induces *Angpt2* transcription in decidualized cells in the PDZ. Taken together, our study illuminates a major responsibility for endocannabinoid signaling in modulating vascular remodeling and integrity. This finding was exemplified in increased vascular leakage at *Cnr1*^*-/-*^*Cnr2*^*-/-*^ implantation sites (6). In conclusion, this investigation reveals that CB1 actions in decidual cells, combined with angiogenic activities driven by CB2, facilitate the formation of the avascular PDZ, which is critical for pregnancy success. Whether other signaling pathways join in this orchestration remains to be seen. Decidualization is a complex process. Although there are unique differences, some common features of this process are evident in mice and humans. As such, decidualization in mice begins with implantation, whereas decidualization in humans occurs in each menstrual cycle, but becomes more intense like in mice, after implantation (1). Whether the PDZ is formed in human implantation or in other subhuman primates is not known. However, the epithelial plaque formed during early pregnancy in macaque may have a similar function (34).

## Materials and Methods

### Animals and Treatments

*Cnr1*^*-/-*^ and *Cnr2*^*-/-*^ mice were generated as described (35,36) and *Cnr1*^*-/-*^*Cnr2*^*-/-*^ females were generated by crossing *Cnr1*^*-/-*^ and *Cnr2*^*-/-*^ mice. All genetically modified mice and WT controls were maintained on a C57BL6 mixed background and were housed in the animal care facility at the Cincinnati Children’s Hospital Medical Center according to NIH and institutional guidelines for laboratory animals. All protocols of the present study were approved by the Cincinnati Children’s Hospital Research Foundation Institutional Animal Care and Use Committee. All mice were housed in wall-mount negative airflow polycarbonate cages with corn cob bedding. They were provided ad libitum with double distilled autoclaved water and rodent diet (LabDiet 5010).

Female mice were mated with WT fertile males to induce pregnancy (vaginal plug=day 1 of pregnancy). Implantation sites were visualized by an intravenous injection of 0.1 mL of 1% Chicago blue dye solution in saline 4 minutes before killing, and the number of implantation sites, demarcated by distinct blue bands, was recorded. Mice were euthanized by cervical dislocation right before tissue collection under deep anesthesia. Mice were killed on different days of pregnancy in accordance to experimental design. i.e day 4, day 6, and day 8 of pregnancy. In each experiment, three to four animals were used to validate the results.

### Fluorescence *in situ* hybridization

*In situ* hybridization was performed as previously described (3). In brief, samples from 3 individual animals in each experimental group were collected. Frozen sections (12 μm) were mounted onto poly-L-lysine-coated slides and fixed in 4% paraformaldehyde in PBS. Following acetylation and permeabilization, slides were hybridized with the DIG-labeled *Bmp2, Ptgs2, Hoxa10, Hif1α, Hif2α, Angpt1, Angpt2* and *Vegfa* probes at 55°C overnight. After hybridization, slides were then washed, quenched in H2O2 (3%), and blocked in blocking buffer (1%). Anti-Dig-peroxidase was applied onto hybridized slides and color was developed by Tyramide signal amplification (TSA) Fluorescein according to the manufacturer’s instructions (PerkinElmer).

### Immunofluorescence

Antibodies for Cytokeratin 8 (Iowa hybridoma bank, 1:100 dilution), CD45 (Biolegend, 1:200 dilution), HIF1α (Cell signaling, 1:200 dilution), HIF2α (Novus, 1:500 dilution), F4/80 (Biolegend, 1:200 dilution) and Scribble (Cell signaling, 1:200 dilution) were used for immunofluorescence staining. All fluorophore conjugated secondary antibodies were from Jackson Immunoresearch. Nuclear staining was performed using Hoechst 33342 (H1399, Molecular Probes, 2µg/ml). Immunofluorescence was performed on fresh-frozen sections. Sections were fixed in 4% paraformaldehyde and washed in PBS. Sections were blocked in 5% BSA in PBS and incubated with primary antibody at 4°C overnight, followed by incubation in secondary antibody for 1 hour in PBS. Immunofluorescence was visualized under a confocal microscope (Nikon Eclipse TE2000).

### Whole-mount immunostaining for 3D imaging

Samples were fixed in Dent’s Fixative (Methanol:DMSO (4:1)) overnight at −20°C. After fixation, tissues were then washed in 100% Methanol three times for 1 h each and bleached with 3% H_2_O_2_ in methanol overnight at 4°C to eliminate pigmentation. The samples were washed in PBS-T containing 0.1% Tween20 three times for 1 h each at room temperature and then blocked in 5% BSA in PBS-T overnight at 4°C. The samples were then incubated with anti-E-cadherin antibody (1:100, 3195 s, Cell Signaling Technology) on a rotor for 7 days at 4°C. After incubation, the samples were then washed in PBS-T 6 times for 1 h each at room temperature and then incubated with Alexa Fluor® 594 AffiniPure Donkey Anti-Rabbit IgG (H + L) (1:300, Jackson ImmunoResearch) on a rotor for 4 days in a light proof tube at 4°C. The samples were stored in the dark until tissue clearing.

### Tissue clearing for 3D imaging

The stained samples were held straight with forceps in 100% methanol for 1 min to align the mesometrial–antimesometrial (M–AM) axis and then dehydrated in 100% methanol for 30 min. Dehydration was followed by tissue clearing by BABB (Benzyl alcohol: Benzyl benzoate (1:2); each reagent from Sigma-Aldrich) for 1 h at room temperature. The samples were then stored in the dark until 3D imaging acquisition (37).

### 3D imaging and processing

3D pictures were acquired by a Nikon multiphoton upright confocal microscope (Nikon A1R) with a 10X objective. To obtain the 3D structure of the tissue, the surface tool Imaris (version 9.2.0., Bitplane) was used (37).

### *In vitro* decidualization of mouse stromal cells

Stromal cells were collected by enzymatic digestion of mouse uteri on day 4 of pregnancy as described previously (21). The cells were cultured in phenol-red free DMEM/F12 medium supplemented with charcoal-stripped 1% FBS (*w*/*v*) overnight prior to the initiation of decidualization by treatment of estradiol (10 nM) and medroxyprogesterone acetate (1 μM) for 6 days. Sample sizes are indicated in each figure.

### *In vitro* decidualization of human uterine fibroblasts

Decidual tissue was dissected from the chorionic layer, and fibroblast cells were isolated and purified to >95% purity by differential plating on plastic as previously described (38). After reaching 90% confluence, the cells were decidualized by the addition of medroxyprogesterone acetate (1 μM), estradiol (E2) (10 nM), and prostaglandin E2 (PGE2) (1 μM) in RPMI medium that contained 2% fetal bovine serum and antibiotics. The medium in each well was changed at 3-day intervals unless indicated otherwise. Sample sizes are indicated in each figure.

### Quantitative RT-PCR

RNA was collected from WT and *Cnr2*^*-/-*^ uterine samples; WT, *Cnr1*^*-/-*^ and *Cnr2*^*-/-*^ primary cells; and human uterine fibroblast cells, with sample sizes indicated in the figures. RNA was analyzed as described previously (39,40). In brief, total RNA was extracted with Trizol (Invitrogen, USA) according to the manufacturer’s protocol.

After DNase treatment (Ambion, USA), 1 µg of total RNA was reverse transcribed with Superscript II (Invitrogen). Real time PCR was performed using primers 5’-ACGTGATGAGGAGGTTCT-3’ (sense) and 5’-AATCTTGCCCAGTTATGC-3’ (anti-sense) for mouse *Prl8a2*; 5’-CATTGGGACTATCTTTGCGG-3’ (sense) and 5’-GGTTCTGGAGAACCTGCTGG-3’ (anti-sense) for mouse *Cnr1;* 5’-GCAGATGTACCGCACTGAGATTC-3’ (sense) and 5’-ACCTTTGGGCTTACTCCATTGATA-3’ (anti-sense) for mouse *Rpl7;* 5’-GCTGCCTAAATCCACTCTGC-3’ (sense) and 5’-TGGACATGAAATGGCAGAAA-3’ (anti-sense) for human *CNR1;* 5’-GATTGGCAGCGTGACTATGA-3’ (sense) and 5’-GATTCCGGAAAAGAGGAAGG-3’ (anti-sense) for human *CNR2;* 5’-CCAAACTGCAACAAGAATG-3’ (sense) and 5’-GTAGACGCACCAGCAGAG-3’ (anti-sense) for human *IGFBP1;* 5’-AAGCTGTAGAGATTGAGGAGCAAA-3’ (sense) and 5’-TCAGGATGAACCTGGCTGACTA-3’ (anti-sense) for human *PRL;* 5’-AAGGAGGAGGGCAGAATCAT-3’ (sense) and 5’-CACACAGGATGGCTTGAAGA-3’ (anti-sense) for human *VEGFA*. 5’-GGACAGCAGGAAAACAGAGC-3’ (sense) and 5’-CACAAGCATCAAACCACCAT-3’ (anti-sense) for human *ANGPT1*. 5’-ATAAGCAGCATCAGCCAACC-3’ (sense) and 5’-AAGTTGGAAGGACCACATGC-3’ (anti-sense) for human *ANGPT2*. 5’-TCATCCAAGAAGCCCTAACG-3’ (sense) and 5’-CGCTTTCTCTGAGCATTCTG-3’ (anti-sense) for human *HIF1α*. 5’-GAACAGCAAGAGCAGGTTCC-3’ (sense) and 5’-GGCAGCAGGTAGGACTCAAA-3’ (anti-sense) for human *HIF2α*. 5’-GAAGGTGAAGGTCGGAGT-3’ (sense) and 5’-GATGGCAACAATATCCACTT-3’ (anti-sense) for human *GAPDH*. Sample sizes are indicated in each figure.

### Statistical Analysis

Data were analyzed by the Student’s *t* tests as indicated in figure legends. Data are shown as mean ± SEM. All *P* values are depicted in the figure legends.

## Acknowledgements

We thank Katie Gerhardt for her efficient editing of the manuscript. This work was supported in parts by NIH grants (DA006668 and HD068524 to SKD).

## Supplemental figure legends

**Supplemental figure 1.**
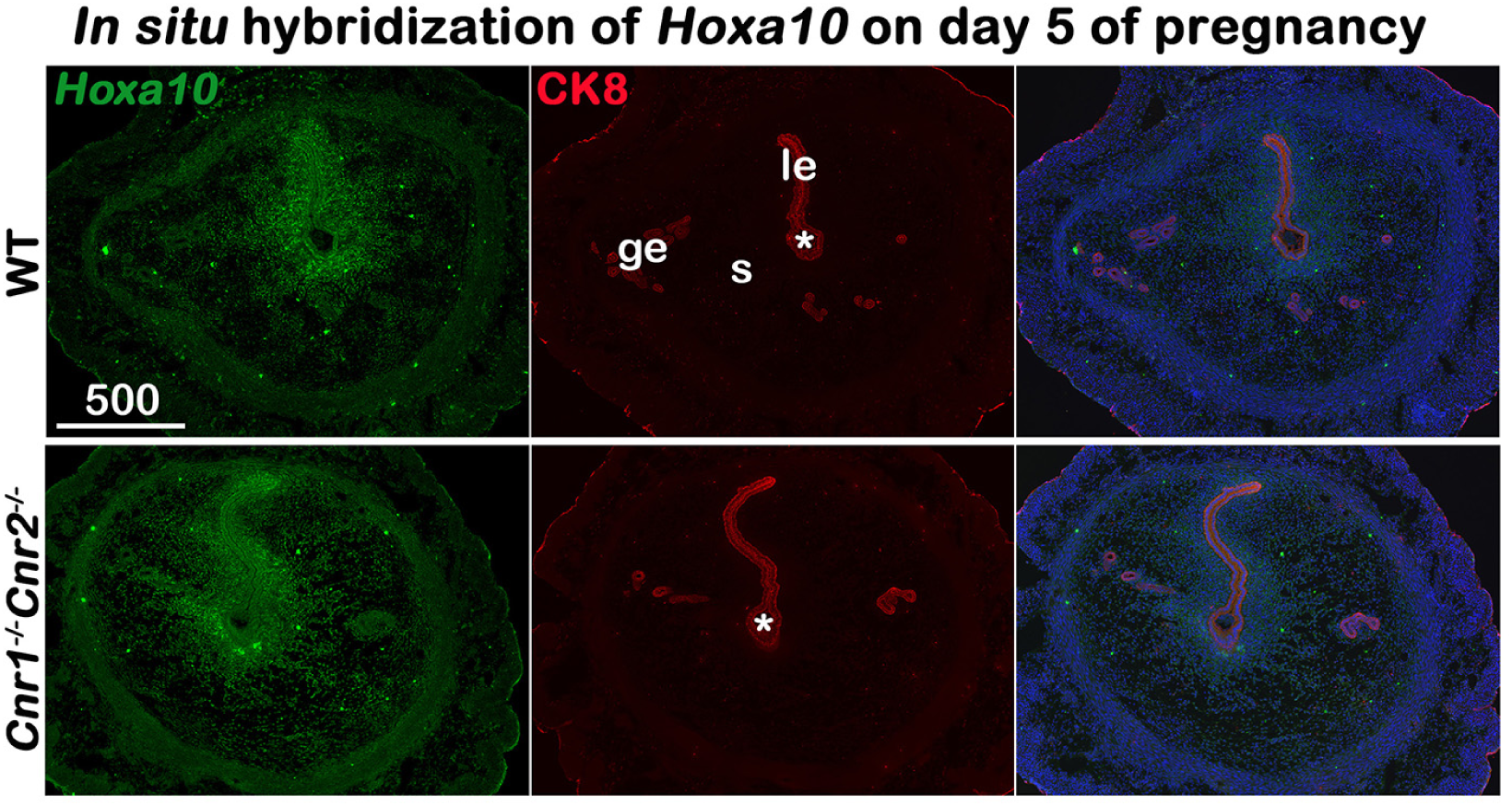
*In situ* hybridization of *Hoxa10* in uteri on day 5 of pregnancy. Decidual responses in *Cnr1*^*-/-*^*Cnr2*^*-/-*^ females are weaker than those in WT females. le, luminal epithelium; s, stroma; ge, glandular epithelium; Asterisks, positions of embryos; Scale bars, 500 μm.

**Supplemental figure 2.**
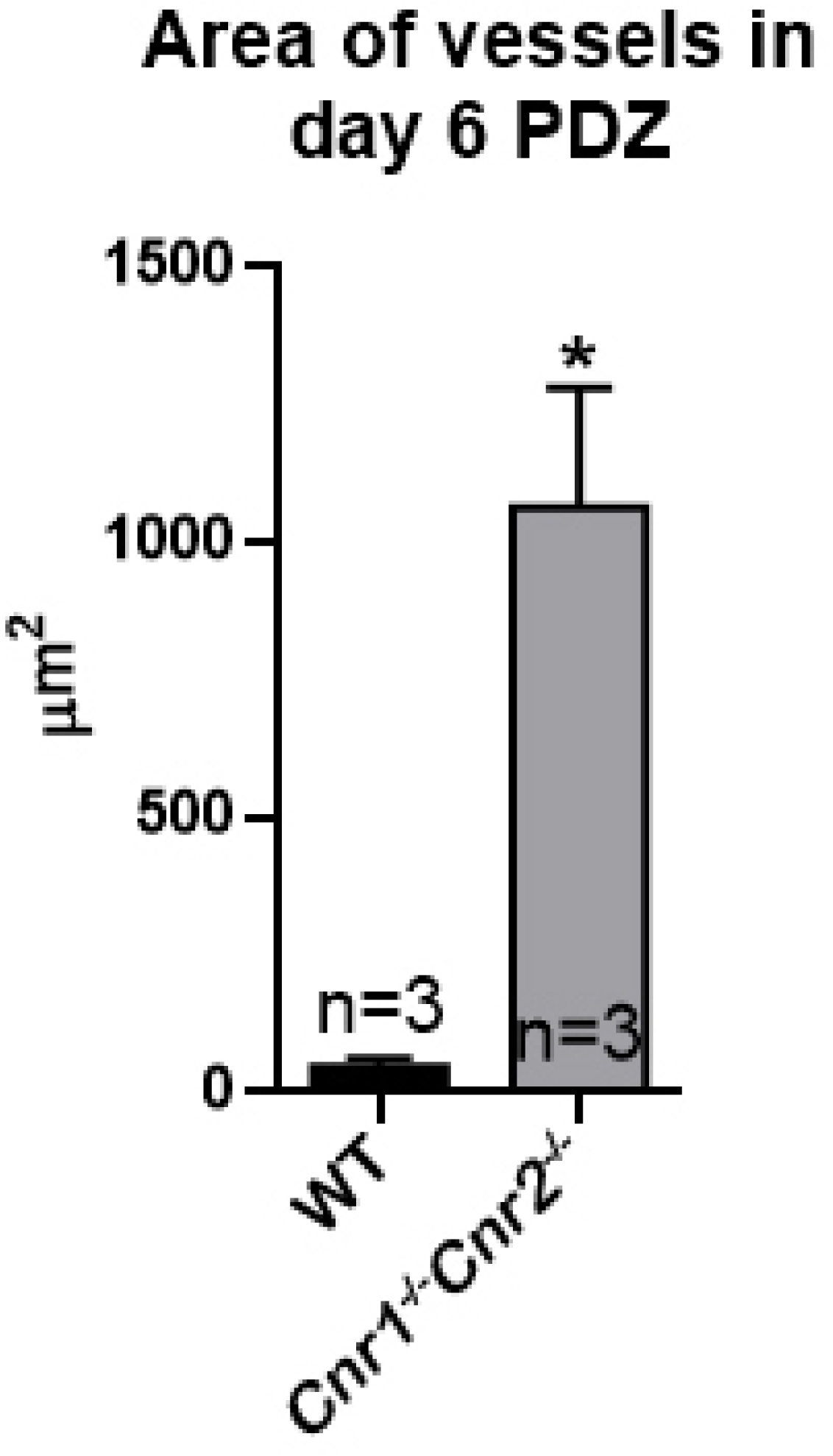
Quantitation of blood vessel area in PDZs on day 6 of pregnancy.

**Supplemental figure 3.**
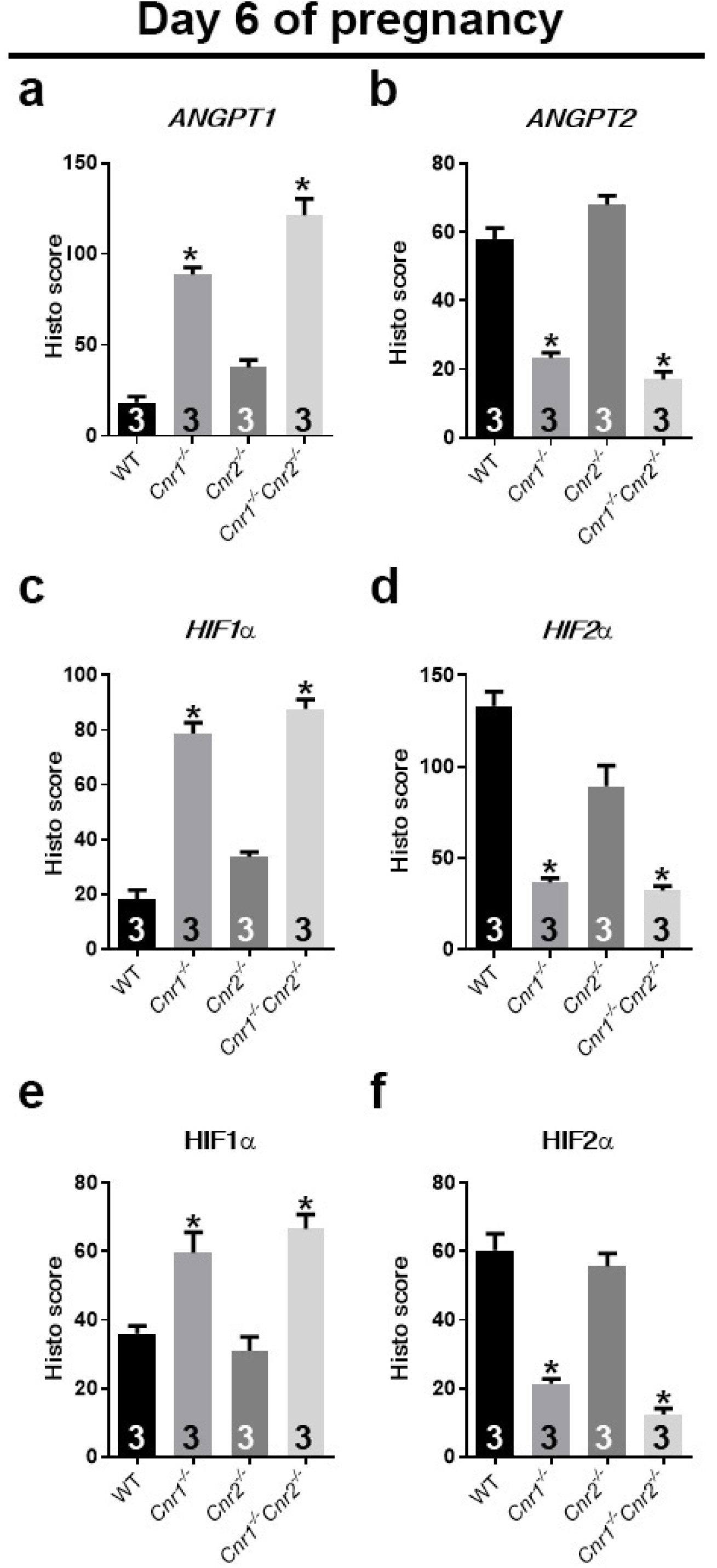
Quantification of signals of angiogenic factors on day 6 of pregnancy. Quantification of *in situ* hybridization signals (a-d) and immunostaining signals (e and f) is plotted. H scores of signals in PDZs are calculated for *ANGPT1* in panel a and *HIF1α* in panels c and e. H scores for *ANGPT2* signals are plotted in panel b. H scores for *HIF2α* signals are plotted in panels d and f. Values are mean ± SEM; Numbers on bars are sample sizes; *P< 0.05, *Student’s t* tests.

**Supplemental figure 4.**
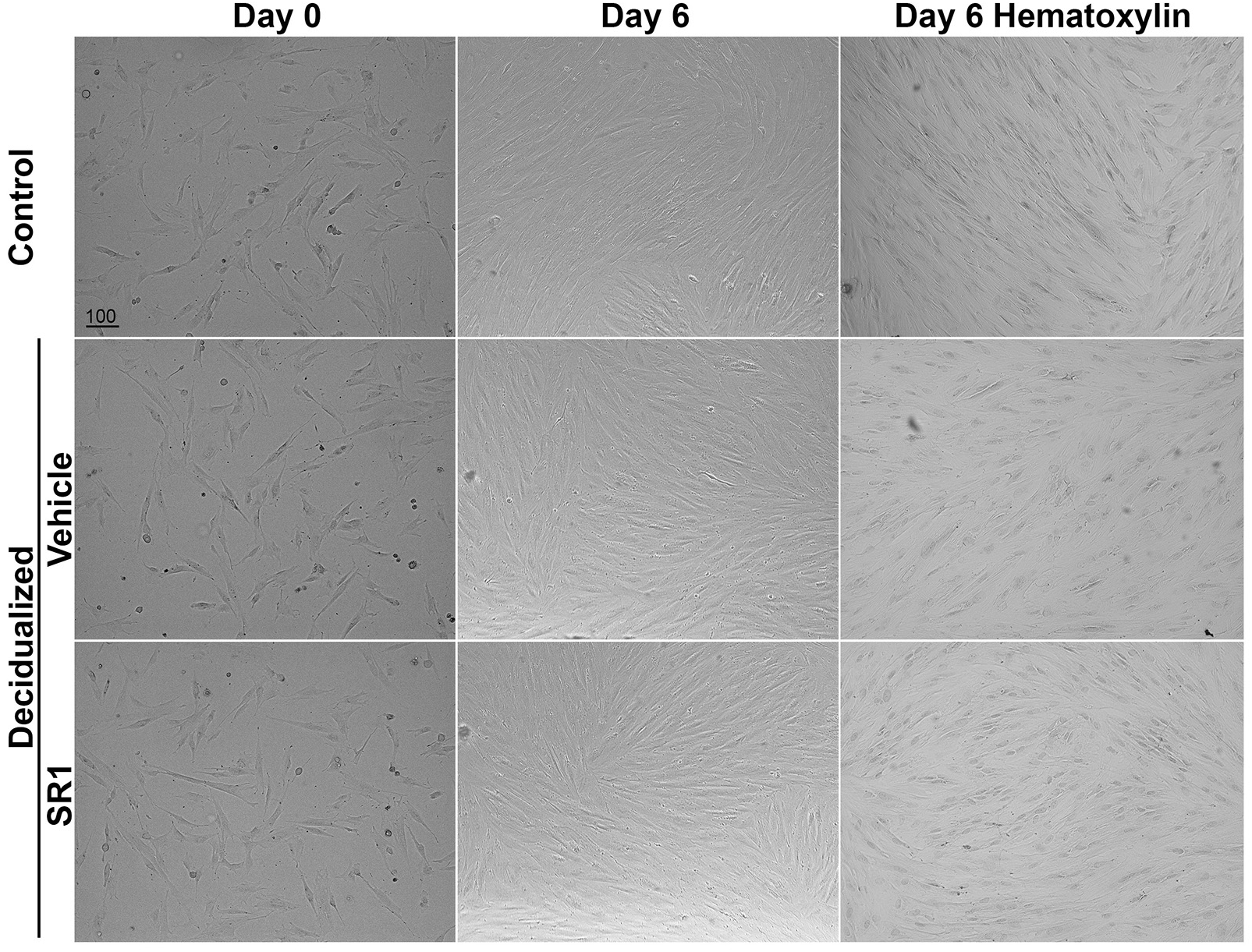
Morphology of Huf cells before and after decidualization. Phase contrast pictures of Huf cells treated with SR1 or vehicle before (day 0) and after (day 6) decidualization were shown. After decidualization, cells were stained with hematoxylin, and pictures were taken after mounting. Scale bars, 100 μm.

**Supplemental figure 5.**
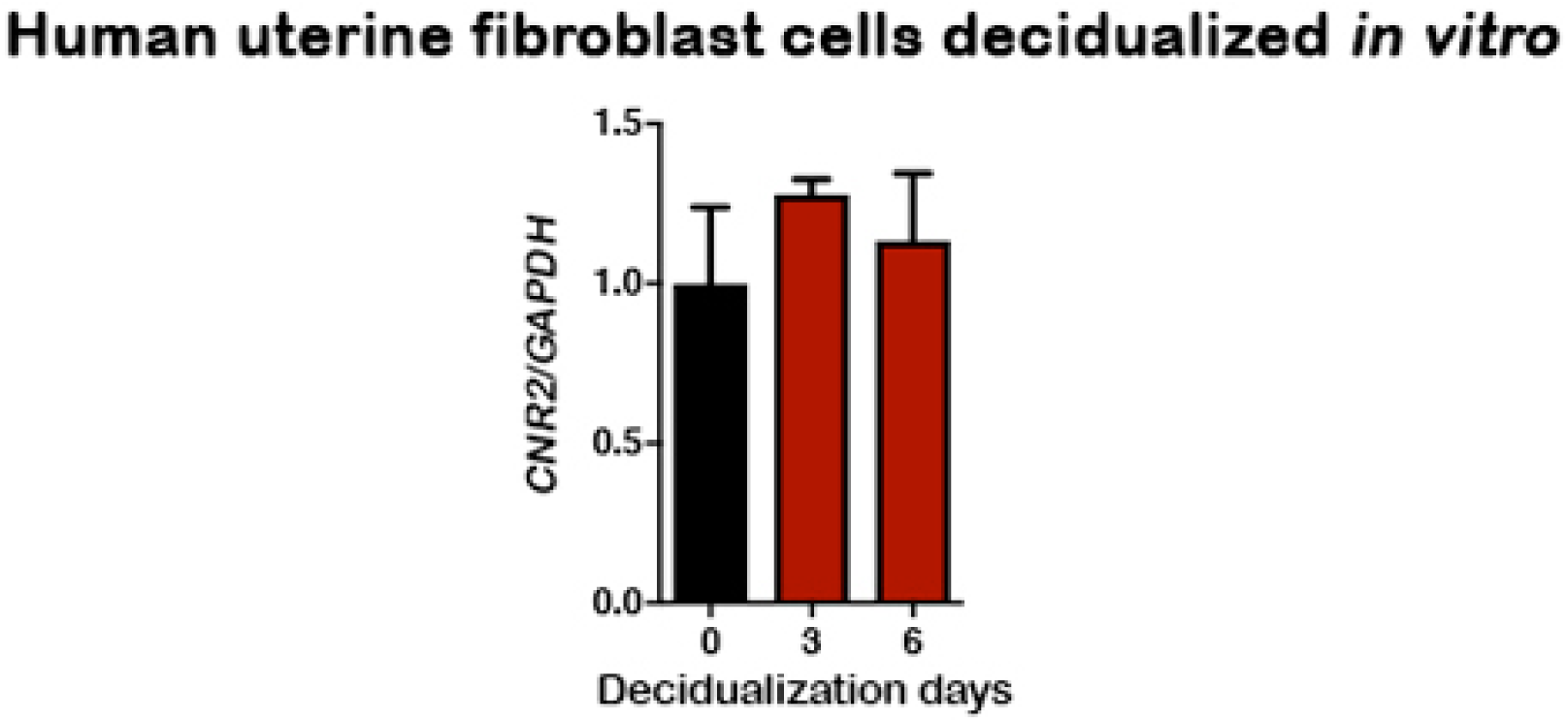
*CNR2* shows no significant change in human uterine fibroblasts during *in vitro* decidualization. Values are mean ± SEM.

